# (-)-Epicatechin modulates skeletal muscle inflammatory response in a mouse model of Limb-Girdle Muscular Dystrophy 2F

**DOI:** 10.64898/2026.05.01.722369

**Authors:** Amairani Cancino-Bello, Mauricio Hernández-Somilleda, Erick Bahena-Culhuac, Estela G. García-González, Oscar Hernández-Hernández, Magally Ramírez-Ramírez, Ramón M. Coral-Vázquez, J. Manuel Hernández-Hernández

**Affiliations:** Laboratorio de Epigenética de la Regeneración Muscular. Departamento de Genética y Biología Molecular. Centro de Investigación sobre el Envejecimiento. Centro de Investigación y de Estudios Avanzados, CINVESTAV-IPN. Ciudad de México. 07360. México; Laboratorio de Medicina Genómica. Departamento de Genética. Instituto Nacional de Rehabilitación - Luis Guillermo Ibarra Ibarra. Ciudad de México. 14389. México; Subdirección de Enseñanza e Investigación, Centro Médico Nacional “20 de Noviembre”, Instituto de Seguridad y Servicios Sociales de los Trabajadores del Estado, Ciudad de México. 03100. Mexico; Sección de Estudios de Posgrado e Investigación, Escuela Superior de Medicina, Instituto Politécnico Nacional, Plan de San Luis y Díaz Mirón s/n, Col. Casco de Santo Tomas, Ciudad de México, México

**Keywords:** RNA-seq, Epicatechin, Limb-Girdle Muscular Dystrophy type 2F, Toll-like receptors, NF-κB2, Interferon-stimulated genes, Skeletal muscle inflammation, innate immunity

## Abstract

Skeletal muscle possesses remarkable regenerative capacity. However, in limb-girdle muscular dystrophy-2F (LGMD2F), this capacity is compromised by persistent innate immune activation, whose transcriptional landscape remains unexplored. In parallel, (-)-Epicatechin has emerged as a promising compound with beneficial effects on muscle and notable anti-inflammatory properties. We therefore used (-)-Epicatechin treatment to test whether it can alleviate LGMD2F-associated transcriptional and immune dysregulation. Here we provide the first transcriptomic characterization of LGMD2F using the *Sgcd^⁻/⁻^* mouse model, along with the first RNA-sequencing-based evaluation of (-)-Epicatechin treatment. We profiled two functionally distinct muscles — the soleus and EDL — through bulk RNA-sequencing coupled with immune cell-deconvolution. *Sgcd^⁻/⁻^* muscles exhibited marked transcriptional dysregulation, more pronounced in the soleus and associated with enhanced innate immune signaling. (-)-Epicatechin induced a muscle- and genotype-dependent transcriptional response: in wild-type animals, the EDL displayed the highest number of differentially expressed transcripts, whereas in *Sgcd^⁻/⁻^* mice, the soleus showed the most prominent response. This shift was accompanied by downregulation of Toll-like receptor and RIG-I-like receptor pathways, along with suppression of NF-κB2 and interferon-stimulated genes. Together, these findings identify innate immune overactivation as a central feature of LGMD2F and reveal (-)-Epicatechin as a context-dependent modulator of muscle-specific transcriptional responses.

## Introduction

Limb-girdle muscular dystrophy type-2F (LGMD-2F) is a debilitating genetic disorder characterized by progressive muscle wasting and weakness, resulting from mutations in the delta-sarcoglycan (δ-SG) gene^[1]^. As an integral component of the Dystrophin-associated protein complex (DAPC), δ-SG deficiency contributes to the loss of membrane integrity resulting in fiber necrosis, calcium overload, oxidative stress, mitochondrial dysfunction and severe cardiomyopathy^[2]^. While the genetic basis of the disease is well-understood, the fundamental question of how differential susceptibility between fast-twitch and slow-twitch fibers, shapes the transcriptional landscape of distinct muscle groups, remains unexplored.

Previous observations have shown that muscle wasting diseases have differential effects on specific muscle groups. In Duchenne Muscular Dystrophy (DMD), fast muscle fibers are preferentially affected^[3]^, as muscle fibers expressing MyHC-2X transcripts disappear early in DMD muscles^[4]^. On the other site, a fiber-type size disproportion, with type-1 muscle fibers being much smaller than type-2 fibers, is frequently seen in congenital myopathies due to mutations in genes such as RYR1, TPM3 and ACTA1. As for sarcoglycanopathies, we recently observed that the slow-twitch muscle soleus contains higher levels of collagen in the connective tissue, in comparison to other muscle groups analyzed on a δ-SG-null mouse^[5]^.

Alongside these observations, a common phenomenon in muscular dystrophies is the presence of chronic inflammation. At the clinical level, Amaro et al. (2025) stratified the severity of LGMD3, the muscular dystrophy caused by α-SG deficiency, based on the cellular composition of the inflammatory infiltrate. By using deconvolution analyses of transcriptional profiles in muscle biopsies, it was observed that the inflammatory profile correlated with the severity of clinical manifestations, allowing to separate the phenotypes from mild to severe. Indeed, pro-inflammatory cytokines and receptors such as CXCL10, CXCL12, CCL5 and CXCR2 were upregulated in severe LGMD3 individuals. Interestingly, an immunological profile in the blood of patients revealed that the percentage of CD8^+^T, CD4^+^TH1T-lymphocytes, and CD69^+^ activated monocytes was higher in LGMDR3 patients compared to control samples^[6]^. However, there is no clear correlation between the severity of inflammation in a given muscle group, with the preferential damage in the fiber type. In line with all these observations, in this study we utilized bulk RNA-sequencing and deconvolution analyses to characterize transcriptional profiles of soleus and EDL in a murine model deficient of the δ-SG protein. In addition, we specifically aimed to define the transcriptional profile of muscle-specific immune infiltration signatures associated with fast- and slow-muscle fibers.

Given that the inflammatory process plays a relevant role in the disease, there is a need to identify natural molecules with anti-inflammatory properties that promote a favorable environment for muscle regeneration in muscular dystrophies. Considering this, (-)-Epicatechin, a flavonoid present in various natural sources^[7–9]^, has shown to be beneficial to muscle homeostasis, while showing promising anti-inflammatory and antioxidant properties in different experimental models^[10–13]^. Hence, we evaluated the capacity of (-)-Epicatechin to mitigate these inflammatory signatures. Our results provide novel insights into the differential responsiveness of slow- and fast-twitch muscles to anti-inflammatory interventions, highlighting the soleus as a primary responder to (-)-Epicatechin-mediated therapeutic effects in the context of LGMD-2F.

## Methods and Materials

### Chemicals

(−)-Epicatechin (Sigma-Aldrich, St. Louis, MO, USA) was prepared as previously described^[14]^

### Animals

Wild-type (WT; B6.129-Sgcd) and δ-sarcoglycan knockout (δ-KO; B6.129-Sgcdtm1Mcn/J) male mice (10–12 weeks) were donated by Dr. Coral-Vázquez. The original strain was obtained from The Jackson Laboratory (Bar Harbor, ME, USA). Genotypes were confirmed using the PCR protocol provided by Jackson Laboratory (strain #004582). At approximately 2.5 months of age, δ-KO mice exhibit a phenotype consistent with human limb-girdle muscular dystrophy (LGMD)^[15]^. Animals were housed in groups of four under controlled environmental conditions (12 h light/dark cycle, 22 ± 2 °C, 50–60% relative humidity). All experimental procedures were conducted in accordance with institutional guidelines (CICUAL, UPEAL) and the Mexican Official Norm (NOM-062-ZOO-1999).

### BaCl₂ injury model and (-)-Epicatechin treatment

Mice were allocated into three groups (n = 3 per group): uninjured, injured vehicle-treated (water + 0.2% DMSO), and injured (-)-Epicatechin-treated. At 10 weeks of age, mice received an intramuscular injection of 1.2% BaCl₂^[16]^. One-hour post-injury, mice were treated by oral gavage twice daily with either vehicle (water + 0.2% DMSO) or 1 mg/kg body weight of (-)-Epicatechin. Mice were euthanized at 2, 4, and 15 days post-injury by cervical dislocation following anesthesia with ketamine (80 mg/kg, i.p.) and xylazine (10 mg/kg, i.p.). For histological analysis, right and left TA muscles were collected and snap-frozen in liquid-nitrogen. Cryosections were stained with Gomori trichrome and images were analyzed using AxioVision SE64 Rel. 4.9.1 software (Carl Zeiss, Jena, Germany). The percentage of damaged area was calculated as the area of granulation tissue divided by the total cross-sectional area.

### Experimental design

The effects of (−)-Epicatechin treatment were evaluated in the soleus (SOL) and extensor digitorum longus (EDL) muscles of δ-KO and wild-type (WT) mice. Four experimental groups were established (n = 6 per group): (a) vehicle-treated WT (WT Ctrl), (b) Epi-treated WT, (c) Epi-treated δ-KO, and (d) vehicle-treated δ-KO (δ-KO Ctrl). Mice received either vehicle (1% DMSO diluted in water) or (-)-Epicatechin (1 mg/kg/body mass) via oral gavage twice daily for 15 consecutive days. On day 16, animals were euthanized by cervical dislocation. The SOL and EDL muscles were dissected and preserved in RNAlater™ (Invitrogen) at −80 °C until further processing for molecular and transcriptomic analyses.

### RNA isolation and Bulk RNA-seq

For each experimental group, Soleus and EDL muscles from *2* mice were pooled to constitute one biological replicate (3 replicates per group). Total RNA was extracted using TRIzol™ reagent, followed by purification with the EZ-10 Spin Column Total RNA Mini-Preps kit (Bio Basic Inc.). To digest genomic DNA, the samples were treated with DNase I (Fermentas). RNA integrity was assessed using an Agilent 2100 Bioanalyzer with an RNA 6000 Nano Chip, and library preparation was performed at LabSerGen (Langebio-CINVESTAV, Mexico) using the MGIEasy RNA Library Prep Kit, NEBNext® Poly(A) mRNA Magnetic Isolation Module, and MGIEasy DNA Adapters-96 Kit. A total of 24 libraries were sequenced on the MGISeq 2000 platform using the DNBSEQ™ technology (DNA nanoballs) to generate ∼17 million single-end 100 bp reads per sample.

### Transcriptomic analyses

The quality of raw sequencing data was assessed using FastQC (v0.11.8) (https://www.bioinformatics.babraham.ac.uk/projects/fastqc/). Reads were aligned to the *Mus musculus* reference genome (Ensembl GRCm39) using the STAR aligner (v2.7.10b)^[17]^. Gene-level read counts were quantified using featureCounts (v2.0.3)^[18]^ by assigning aligned reads to annotated gene features. Genes with 10 or less counts across all samples were filtered prior to analysis. Differential gene expression analysis was performed using DESeq2 (v1.44.0)^[19]^ from raw count matrices. Genes with an adjusted p-value ≤ 0.05 were considered differentially expressed (DEGs); no fold-change threshold was applied. EnhancedVolcano (v1.22.0) was used for volcano plots. The overlap among DEG sets was visualized using UpSetR (v1.4.0) and VennDiagram (v1.7.3). Spearman’s rank correlation analysis of log₂ fold change values was performed to assess the concordance of transcriptional changes between Sgcd⁻/⁻ soleus and EDL muscles for genes identified as DEGs in both tissues. All analyses were conducted in R (v4.4.0), with data manipulation and visualization performed using dplyr (v1.1.4) and ggplot2 (v3.5.1).

### GO and KEGG analysis

Gene Ontology (GO) enrichment analysis was performed on DEGs using the enrichGO function from clusterProfiler (v4.12.6)^[20]^, with org.Mm.eg.db (v3.19.1) as the annotation database. Gene symbols were used as input (keyType = “SYMBOL”), and each GO domain (Biological Process, Cellular Component, and Molecular Function) was analyzed independently. For KEGG pathway enrichment, gene symbols were first mapped to Entrez IDs using the bitr function, and the enrichKEGG function was applied with the organism set to “mmu”. In both analyses, p-values were adjusted using the Benjamini–Hochberg method, with a significance cutoff of adjusted p < 0.05 and q-value < 0.2.

### Immune cell deconvolution analysis

Immune cell composition was estimated from bulk RNA-seq data using two complementary deconvolution approaches to increase the robustness of the results. mMCP-counter^[21]^, implemented through the mMCP-counter R package (v1.1.0), was used to estimate enrichment scores for eight immune and two stromal cell populations based on transcriptomic signatures validated in mouse tissues. quanTIseq^[22]^, implemented through the quantiseqr R package (v1.12.0), was used to estimate relative fractions of ten immune cell populations: B cells, dendritic cells, M1 and M2 macrophages, monocytes, neutrophils, NK cells, CD4⁺ T cells, CD8⁺ T cells, and regulatory T cells (Tregs). The algorithm was run using the TIL10 signature matrix with parameters is_arraydata = FALSE, is_tumordata = FALSE (as the samples correspond to non-tumor skeletal muscle tissue), and scale_mRNA = TRUE to enable mRNA content normalization. The uncharacterized “Other” fraction was excluded from downstream analyses. For both methods, a TPM-normalized gene expression matrix with gene symbols as identifiers (GCRm39 assembly) was used as input. Results were visualized as stacked bar plots using ggplot2 (v4.0.0) in R (v4.4.0).

### cDNA synthesis and qPCR

cDNA was synthesized using M-MLV reverse transcriptase (Invitrogen) and Oligo dT (20 primer) (Invitrogen). Real-time PCR was carried out using the Rotor-Gene Q system (Qiagen) with Kappa Sybr-Fast (Roche #kk4602). Primer sequences for the target genes are provided in Supplementary Table 1. Relative gene expression levels were determined using the comparative Ct-(ΔΔCt) method, with normalization to the TATA-box binding protein (TBP) gene.

### Statistical analysis

Statistical analyses were performed using R (v4.4.0) and GraphPad Prism (v10.3.0). Data are presented as mean ± SD. For qPCR data, comparisons between two groups were performed using a two-tailed unpaired Student’s t-test. For VST-normalized expression values, comparisons between two groups were performed using a two-tailed unpaired Student’s t-test, while comparisons among three or more groups were analyzed by one-way or two-way ANOVA, as appropriate, followed by Tukey’s (for comparisons among all groups) or Šídák’s (for pre-selected pairwise comparisons) post-hoc tests. Statistical significance was set at p< 0.05 and is indicated as *p< 0.05, **p< 0.01, ***p< 0.001.

## Results

### δ-sarcoglycan deficiency upregulates innate immune and interferon-stimulated gene programs in *Sgcd^−/-^* mice

To investigate transcriptional changes contributing to muscle degeneration in a murine model of limb-girdle muscular dystrophy 2F, we performed bulk RNA-sequencing on soleus and EDL from δ-SG-null and wild-type mice (Figure 1A). Principal component analysis revealed that muscle type is the primary source of transcriptional variation (PC1), whereas genotype-associated differences were evident in PC2 (Figure 1B). Thus, the transcriptional profiles of wild-type muscles confirmed known expression signatures distinguishing oxidative and glycolytic muscle groups. Differential expression analysis of wild-type muscles identified 2,205 genes upregulated in EDL and 2,278 genes upregulated in soleus (Figure 1C, Supplementary list 1). After gene ontology enrichment analysis, upregulated genes in the soleus were enriched in pathways such as fatty acid metabolism, fatty acid oxidation, and mitochondrial protein-containing complexes. In contrast, EDL showed specific gene expression in pathways including glycolysis, contractile fiber, muscle cell differentiation, and muscle cell development (Figure 1D, Supplementary list 2). Further differential gene expression analysis between dystrophic and wild-type mice revealed a total of 1,958 upregulated and 1,512 downregulated genes in the δ-SG-null model (Supplementary Figure 1A, Supplementary list 12). GO enrichment and KEGG analysis of soleus and EDL combined upregulated genes were predominantly associated with immune response pathways, including innate immune response activation, NF-kappa B signaling, Toll-like receptor signaling, chemokine signaling, and inflammasome-mediated signaling pathway (Supplementary Figure 1B and D, supplementary List 13-14)^[23–27]^. Centplot of GO enrichment results highlighted key inflammatory mediators within these pathways, including Oas3, Oas1a, Oasl1, Ly96, Myd88, Cd68, Serpinb9, Isg15, Serpine1, Tgfbr2, Tnf, Tlr2, and Anxa family of genes^[28–30]^ (Supplementary Figure 1C). Notable among these were Tnf, Cd68, Isg15, and Tlr2, reflecting the chronic inflammatory state, characteristic of muscular dystrophy^[31]^. Conversely, GO enrichment and KEGG analysis of downregulated genes in both muscles were enriched in pathways essential for muscle function, including muscle cell differentiation, generation of precursor metabolites and energy, MAPK and Wnt signaling, and cellular respiration^[32,33]^ (Supplementary Figure 2A and 2C, Supplementary lists 15 and 16). Representative genes included mTOR, Ndufs1, Nnt, Pdha1, Oxct1, Cox5a, ATP5PB, Trim32, Acta1, Sirt3, Sirt4, and Sirt5. Together, these alterations indicate impaired metabolic capacity and regenerative potential in δ-SG-null muscles, consistent with a characteristic transcriptional signature of muscular dystrophy. While inflammatory pathways become hyperactivated as a response to ongoing muscle damage, the fundamental cellular machinery required for energy production, muscle contraction, and tissue regeneration becomes progressively impaired. Interestingly, transcriptional identity of muscle groups was preserved in δ-SG-null mice, with only reduction in expression levels but no major shift in fiber-type-defining genes. Consistently, key EDL-specific markers (e.g., Myh4, Tnnc2, Pvalb, Stat5b, and Wnt4)^[34]^ remained highly expressed in EDL, whereas soleus-specific markers (e.g., Myh7, Myh2, Pdlim1, Bdh1, and Lincmd1)^[35]^ were also maintained (Figure 1E). VST-normalized expression values for fiber-type marker genes in EDL^[34]^ such as Ppl7 (Net39), Stat5b, and Wnt4 remained upregulated, while soleus-specific markers, including Pdlim1, Lincmd1 and Bdh1 maintained their preferential expression in soleus, regardless of the genotype (Figure 1F). Thus, we observed changes in expression levels of functional genes as a consequence of muscular dystrophy, without evident compromise of muscle-type identity.

**Figure 1.**
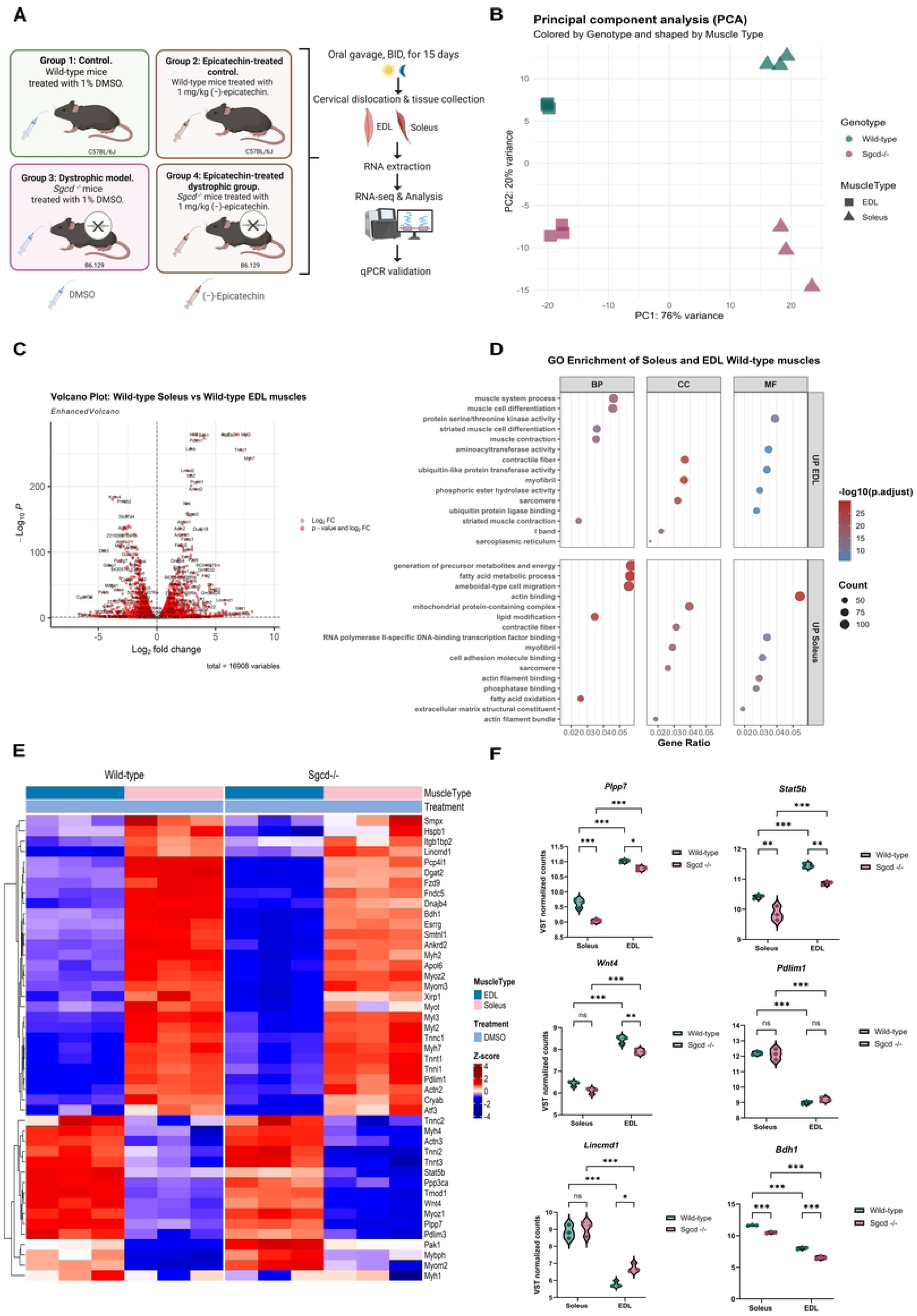
Differential transcriptomic analysis of soleus and extensor digitorum longus (EDL) muscle samples from wild-type mice. **A)** Schematic of experimental design, group assignments, and treatments. **B)** Principal component analysis (PCA) of the merged datasets analyzed in this study shows no evidence of batch effects. **C)** Volcano plot comparing gene expression between soleus and extensor digitorum longus (EDL) muscles from wild-type mice. Each point represents an individual gene. Genes with significant differential expression (p.adj < 0.05) are colored, while non-significant genes (p.adj ≥ 0.05) are shown in gray. n = 3 biological replicates per group. **D)** GO enrichment analysis of soleus and EDL wild-type muscles. Differentially expressed genes were identified using DESeq2, with upregulated genes defined as those with log2FC > 0 and p.adj ≤ 0.05. Biological Process (BP), Cellular Component (CC), and Molecular Function (MF) terms are listed for genes upregulated in each tissue. GO enrichment analysis was performed using clusterProfiler with significance thresholds of p.adj < 0.05 and q-value < 0.2. **E)** Heatmap of 45 fiber-type-specific marker genes in wild-type and *Sgcd*⁻/⁻ EDL and Soleus muscles. Color scale represents VST-normalized expression values scaled by Z-score. Hierarchical clustering was applied to rows. **F)** Violin plots displaying the distribution of VST-normalized expression values for fiber-type marker genes in soleus and EDL muscles from wild-type and *Sgcd⁻/⁻* mice. Two-way ANOVA was performed, followed by Tukey’s post hoc test. Asterisks indicate statistical significance: p < 0.05 (\**), p < 0.01 (**), p < 0.001 (\*\*\**).

**Figure 2.**
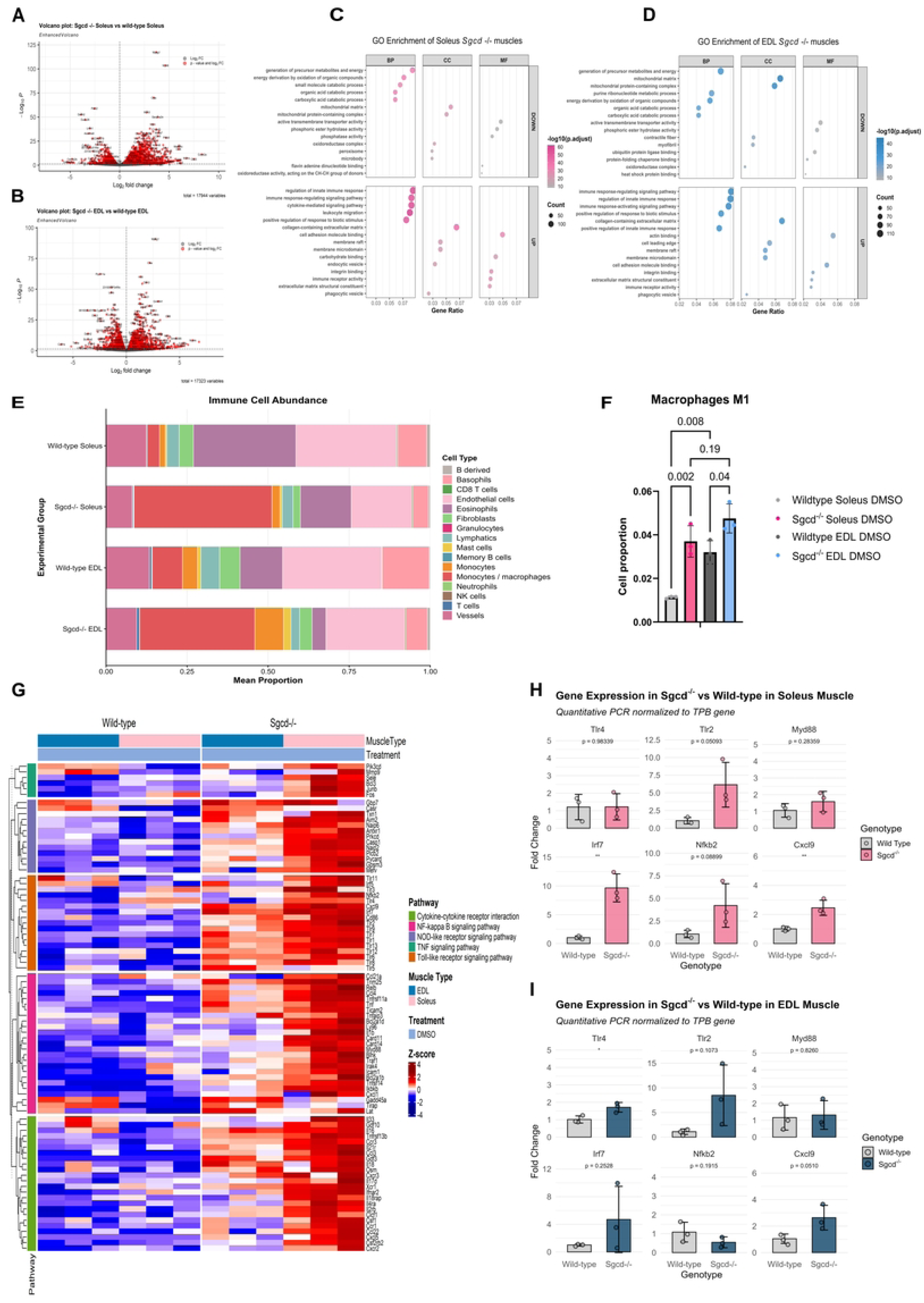
Impact of Sgcd deficiency on Soleus and EDL muscle gene expression. **A)** and **B)** Volcano plots comparing gene expression between Sgcd^−/-^ muscles versus Wild-type. Each point represents an individual gene. Red points indicate significantly differentially expressed genes (p.adj ≤ 0.05), while gray points represent non-significant genes. Differentially expressed genes were identified using DESeq2. n = 3 biological replicates per group*⁻*. **C)** and **D)** Gene Ontology (GO) enrichment analysis of differentially expressed genes in *Sgcd⁻/⁻* soleus and EDL muscles. Upregulated genes were defined as those with log2FC > 0 and p.adj ≤ 0.05, while downregulated genes were defined as those with log2FC < 0 and p.adj ≤ 0.05. Biological Process (BP), Cellular Component (CC), and Molecular Function (MF) terms are shown for both upregulated and downregulated genes. GO enrichment analysis was performed using clusterProfiler with significance thresholds of p.adj < 0.05 and q-value < 0.2. **E)** Bar plot showing estimated immune cell abundance in EDL and Soleus muscles from *Sgcd*⁻/⁻ and wild-type mice, as determined by mMCP-counter deconvolution. Monocytes/macrophages were the most abundant immune cell population in *Sgcd*⁻/⁻ muscles relative to wild-type controls (n = 3 per group). **F)** Estimated abundance of M1 macrophages in EDL and Soleus muscles from *Sgcd*⁻/⁻ and wild-type mice, as determined by quanTIseq deconvolution. Data are presented as mean ± SD (n = 3 per group). One-way ANOVA followed by Šídák’s post hoc test; *p < 0.05, **p < 0.01, ***p < 0.001. **G)** Heatmap of immune response-related genes from NF-κB signaling, NOD-like receptor signaling, TNF signaling, and Toll-like receptor signaling pathways in wild-type and *Sgcd*⁻/⁻ EDL and Soleus muscles. Color scale represents VST-normalized expression values scaled by Z-score. Rows represent genes (n = 123) and columns represent individual samples. Hierarchical clustering was applied to rows. **H)** and **I)** RT-qPCR validation of six immune-related genes (*Tlr2*, *Tlr4*, *Myd88*, *Irf7*, *Nfkb2*, *Cxcl9*) in EDL and Soleus muscles from *Sgcd*⁻/⁻ and wild-type mice. Expression levels were normalized to *Tbp* and presented as fold-change relative to wild-type controls. Data represent mean ± SD (n = 3 biological replicates per group). Two-tailed unpaired Student’s t-test: *p < 0.05, **p < 0.01, ***p < 0.001.

### Soleus and EDL muscles exhibit divergent transcriptional responses to δ-sarcoglycan deficiency while preserving tissue-specific gene expression signatures

Given that muscle-type identity is maintained in *Sgcd^⁻/⁻^* mice, we next sought to characterize the disease-specific transcriptional changes by comparing gene expression profiles between *Sgcd^⁻/⁻^* and wild-type muscles. It is well established that specific muscle groups exhibit differential susceptibility to damage in muscular dystrophies^[3,36,37]^. Then, we sought to define this specific gene expression profile in our LGMD-2F model. First, we analyzed transcriptional profiles of fast-twitch and slow-twitch muscles in response to δ-sarcoglycan deficiency, relative to their respective wild-type controls. We identified 1,786 upregulated and 1,193 downregulated genes in the *Sgcd⁻/⁻* soleus, whereas 1,457 upregulated and 1,596 downregulated genes in the *Sgcd⁻/⁻* EDL (Figures 2A-B, supplementary lists 3-4). From this, 477 unique genes were downregulated in soleus and 876 in EDL in the KO mouse, whereas 713 genes were downregulated in both muscles. In addition, we identified 802 unique genes upregulated in soleus and 477 unique upregulated genes in EDL, whereas 977 genes were increased in both muscles, as shown in the upset plot (Supplementary figure 3A). Consistent with this, gene ontology analysis of genes overexpressed in the Sgcd⁻/⁻ soleus and EDL revealed very similar enrichment of pathways such as “regulation of innate immune response”, “immune response-signaling pathway”, “cytokine-mediated signaling pathway” and “leukocyte migration”. As for downregulated genes, biological processes showed pathways involved in “generation of precursor metabolism and energy”, “mitochondrial matrix”, “oxidation of organic compounds” and “organic acid catabolic processes” (Figures 2C and 2D; Supplementary List 5-6).

To evaluate the degree of transcriptional similarity between muscles deficient of δ-SG, we compared the log2 fold-change values of the 1,700 DEGs shared between *Sgcd⁻/⁻* soleus and *Sgcd⁻/⁻* EDL, as identified by the UpSet plot analysis (Supplementary Figure 3A). The resulting scatter plot (Supplementary Figure 3B) revealed a strong positive Spearman-rank correlation (ρ = 0.935, p < 2.2 × 10⁻¹⁶), indicating that the transcriptional response to δ-sarcoglycan loss is highly conserved between both muscle types. Notably, soleus consistently exhibited larger fold-change magnitudes than the EDL. Having in mind our observations in soleus, we performed RT-qPCR analysis of selected immune-related transcripts (*Tlr4, Tlr2, Myd88, Nfkb2, Irf7,* and *Cxcl9*) in both *Sgcd⁻/⁻* mice muscles (Supplementary Figure 3C). Consistently, all six genes showed higher expression levels in the soleus compared to the EDL, supporting the enhanced inflammatory transcriptional response in this tissue.

Inflammation plays fundamental roles in the development of muscular dystrophies and in such context, it severely affects muscle regeneration^[38–40]^. Considering this, we analyzed the relative immune infiltration status in soleus and EDL from δ-SG-null and control mice. Thus, deconvolution analyses were conducted using mMCP-counter^[6]^ based on the expression profiles of each muscle. This allowed us to obtain the relative abundance of inflammatory infiltrate cells in each sample. The proportion of basophils was reduced in both muscles from the KO mice, alongside eosinophils, fibroblasts, and vessels — with the soleus showing a more pronounced reduction than the EDL — whereas the proportion of monocytes and macrophages was increased in both muscles of the dystrophic mice (Figure 2E). Again, this increase was more robust in soleus than in EDL. Given that macrophages play a central role in the resolution of inflammation through dynamic shifts in their activation states—from predominantly pro-inflammatory (M1-like) toward anti-inflammatory and pro-resolving (M2-like) phenotypes^[41]^—we subsequently applied quanTIseq^[22]^ to track the M1-like macrophages profiles. Indeed, M1-like macrophages in δ-SG-null were significantly more abundant in soleus (P=0.002) than in EDL (P=0.04) compared to the wild type (Figure 2F, supplementary figure 4A). These results suggest that persistent inflammation mediated by M1-like macrophages exacerbates the dystrophic phenotype, and that soleus may be more prone to chronic inflammation and damage than EDL. Consistent with these, KEGG pathway enrichment analysis, restricted to the 908 upregulated genes exclusive to the Sgcd⁻/⁻ soleus, revealed significant enrichment of inflammatory and immune-related pathways, including Cytokine–cytokine receptor interaction, NF-κB, NOD-like receptor, TNF, and Toll-like receptor signaling, as shown in the heatmap in Figure 2G (supplementary List 7). As for the soleus, RT-qPCR confirmed the upregulation of multiple immune-related genes, including Tlr2, Nfkb2, Cxcl9, and Irf7 (Figure 2H), as well as Rigi, Lgp2, Ifih1, Ifit3, Ifit1, Ddx60, Mx1, Isg15, Oas3, Oasl2, Oasl1, and Oas1a (Supplementary Figure 4B). In contrast, although EDL also showed enrichment of immune-related pathways such as activation of innate immune response, regulation of leukocyte activation, and T cell activation, the number of genes within these categories was lower than in soleus. Consistently, RT-qPCR analysis in EDL revealed no significant differences for Myd88, Irf7, and Nfkb2 between δ-SG-null and wild-type mice (Figure 2I), in contrast to the robust changes observed in the soleus, although *Tlr4* was significantly increased.

Together, these results indicate that soleus mounts a stronger and more sustained inflammatory response, likely contributing to the greater susceptibility of slow-twitch fibers to damage compared to fast-twitch fibers.

### The transcriptional effects of (−)-Epicatechin are shaped by muscle-type and genotype

We have observed that (-)-Epicatechin administration after BaCl₂-induced damage in wild-type mice, rapidly reduced cellular infiltrate as well as the area of muscle damage in tibialis anterior (Supplementary Figure 5A and 5B). Consistent with these findings, previous studies have reported that (−)-Epicatechin enhances mitochondrial function, improves oxidative capacity, and promotes metabolic adaptations in skeletal muscle across different physiological contexts^[42–46]^. Thus, knowing that the inflammation may explain some of the differences in the transcriptional response between soleus and EDL of the δ-SG-null mice, we analyzed gene expression changes after (-)-Epicatechin administration (Figure 1A). As seen in figure 3A, muscle groups and genotypes were the main source of transcriptional differences, and (-)-Epicatechin administration had no evident effects on this distribution. Interestingly, our results revealed that transcriptional markers of immune infiltration decreased in response to (-)-Epicatechin in *Sgcd^−/-^* mice, again with no evident changes on muscle-type defining genes. In soleus, downregulation of 101 genes was observed (Figure 3B, supplementary list 8), whereas only 38 genes were reduced in EDL (Figure 4A, supplementary list 10), we also found that (-)-Epicatechin upregulated a limited number of genes in both muscles (38 in Soleus and 20 in EDL); nevertheless, these genes were associated with muscle contraction-related pathways, including muscle system process and skeletal muscle contraction (Figure 3C). Gene ontology analysis of differentially expressed genes in the *Sgcd^−/-^* mice treated with (-)-Epicatechin showed downregulation of pathways involved in positive regulation of innate immune response, response to interferon beta and antiviral response. Interestingly, genes involved in pathways such as acute inflammatory response, canonical NF-KappaB signal transduction, cytokine production involved in immune response and positive regulation of innate immune response, were downregulated to levels comparable to those observed in DMSO-treated WT mice (supplementary lists 9 and 11). Consistently, pathways related to the regulation of AIM2 inflammasome complex and regulation of type-I interferon production, were also reduced (Figure 3D). In addition, quantitative-PCR analyses confirmed a significant downregulation of key immune genes including Tlr2, Irf7, Nfkb2, Cxcl9, Ddx58, Ifit1, Mx1, Isg15, Oas3, and Oasl2. Additionally, we observed trends toward downregulation in Tlr4, Myd88, Dhx58, Ifit3, Ddx60, Oasl1, and Oas1a, although these changes were not statistically significant (Figure 3E).

**Figure 3.**
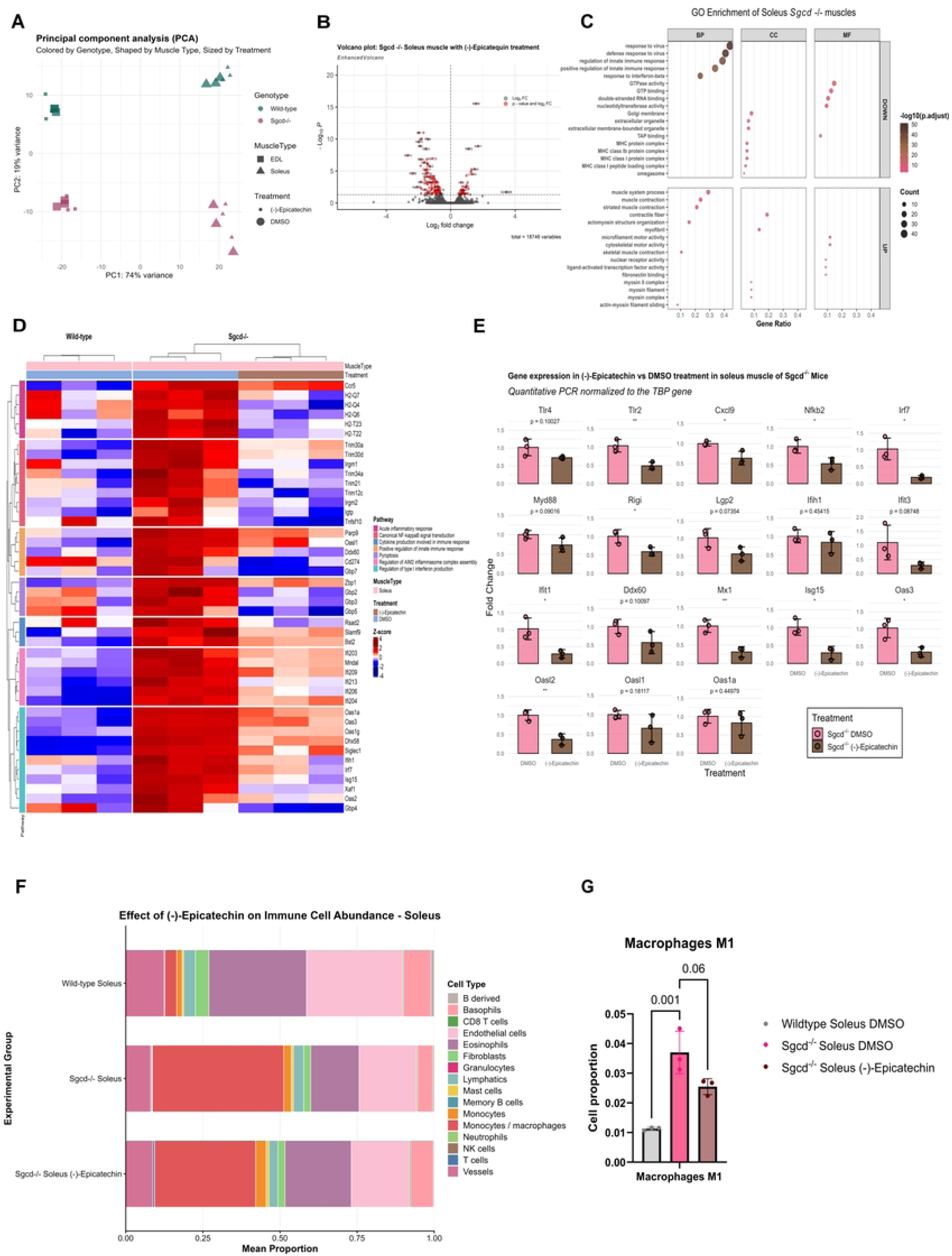
Epicatechin attenuates inflammatory gene expression in *Sgcd*⁻/⁻ Soleus muscle. **A)** Principal component analysis (PCA) of the merged datasets analyzed in this study shows no evidence of batch effects. **B)** Volcano plot of differentially expressed genes in *Sgcd*⁻/⁻ Soleus muscle comparing epicatechin-treated versus vehicle-treated (DMSO) mice. The dashed line indicates the significance threshold at adjusted p-value < 0.05. Significant genes (red) are distributed as upregulated (right) or downregulated (left) relative to the fold-change axis. **C)** Gene Ontology (GO) enrichment analysis of differentially expressed genes in *Sgcd⁻/⁻* soleus muscle after epicatechin treatment. Upregulated genes were defined as those with log2FC > 0 and p.adj ≤ 0.05, while downregulated genes were defined as those with log2FC < 0 and p.adj ≤ 0.05. Biological Process (BP), Cellular Component (CC), and Molecular Function (MF) terms are shown for both upregulated and downregulated genes. GO enrichment analysis was performed using clusterProfiler with significance thresholds of p.adj < 0.05 and q-value < 0.2. **D)** Heatmap of 44 differentially expressed genes (adjusted p-value ≤ 0.05) in *Sgcd*⁻/⁻ and wild-type Soleus muscle associated with Acute inflammatory response, Canonical NF-κB signal transduction, Pyroptosis, and Regulation of AIM2 inflammasome complex assembly pathways. Color scale represents VST-normalized expression values scaled by Z-score. Hierarchical clustering was applied to both rows and columns. **E)** RT-qPCR validation of 18 immune-related genes in *Sgcd*⁻/⁻ Soleus muscle comparing epicatechin-treated versus vehicle-treated (DMSO) groups. Expression levels were normalized to *Tbp* and presented as fold-change relative to the vehicle-treated group. Data represent mean ± SD (n = 3 biological replicates per group). Statistical significance was determined using two-tailed unpaired Student’s t-test: *p < 0.05, **p < 0.01, ***p < 0.001. **F)** Bar plot of estimated immune cell abundance in Soleus muscles from *Sgcd*⁻/⁻ vehicle-treated (DMSO), *Sgcd*⁻/⁻ epicatechin-treated, and wild-type mice, as determined by mMCP-counter deconvolution (n = 3 per group). **G)** Estimated abundance of M1 macrophages in Soleus muscles from *Sgcd*⁻/⁻ vehicle-treated (DMSO), *Sgcd*⁻/⁻ epicatechin-treated, and wild-type mice, as determined by quanTIseq deconvolution. Data are presented as mean ± SD (n = 3 per group). One-way ANOVA followed by Šídák’s post hoc test: *p < 0.05, **p < 0.01, ***p < 0.001.

**Figure 4.**
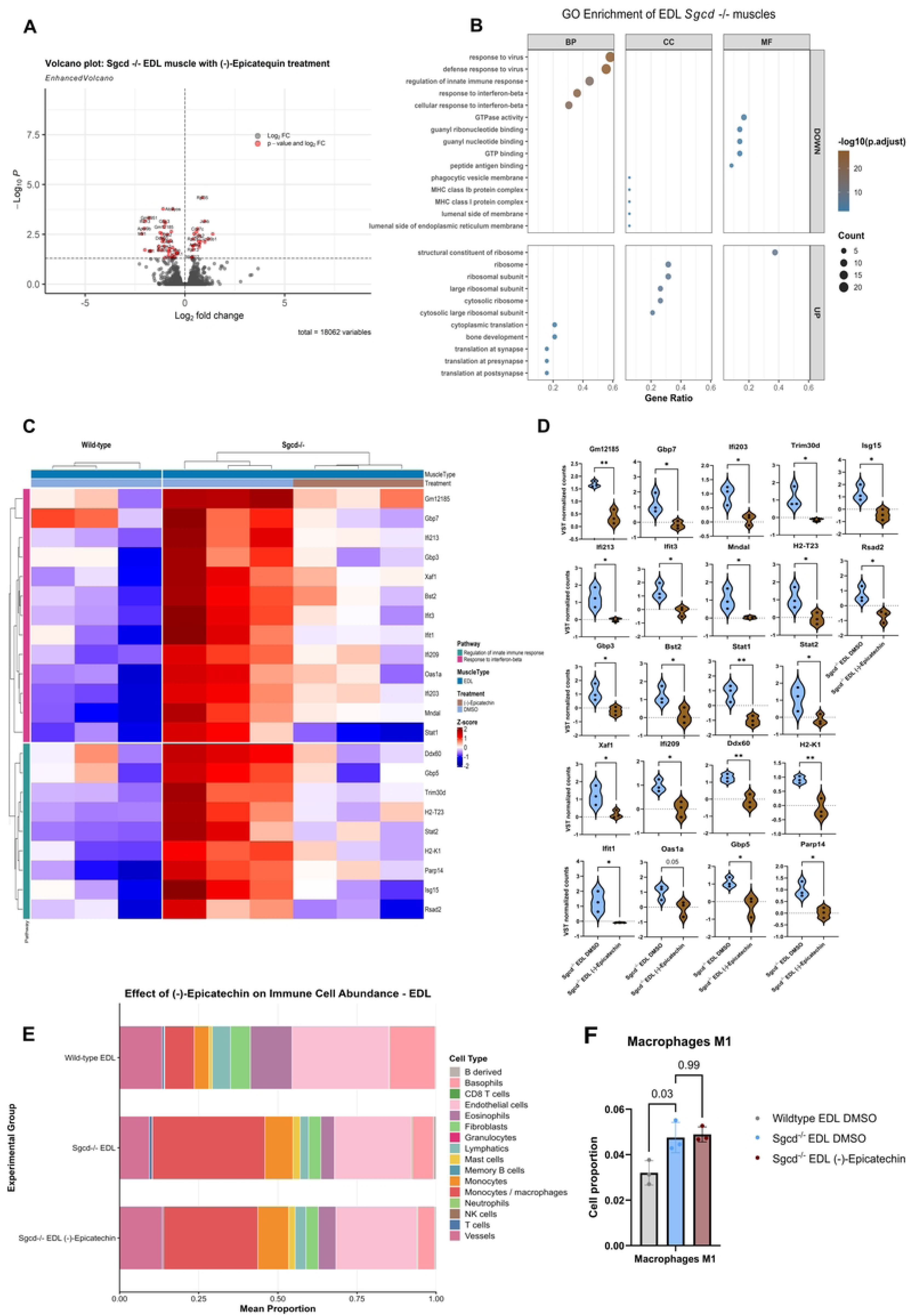
Epicatechin shows limited transcriptional effects in Sgcd-/- EDL muscle. **A)** Volcano plot of differentially expressed genes in *Sgcd*⁻/⁻ EDL muscle comparing epicatechin-treated versus vehicle-treated (DMSO) mice. The dashed line indicates the significance threshold at adjusted p-value < 0.05. Significant genes (red) are distributed as upregulated (right) or downregulated (left) relative to the fold-change axis. **B)** Gene Ontology (GO) enrichment analysis of differentially expressed genes in *Sgcd⁻/⁻* EDL muscle after epicatechin treatment. Upregulated genes were defined as those with log2FC > 0 and p.adj ≤ 0.05, while downregulated genes were defined as those with log2FC < 0 and p.adj ≤ 0.05. Biological Process (BP), Cellular Component (CC), and Molecular Function (MF) terms are shown for both upregulated and downregulated genes. GO enrichment analysis was performed using clusterProfiler with significance thresholds of p.adj < 0.05 and q-value < 0.2. **C)** Heatmap of 22 differentially expressed genes (adjusted p-value ≤ 0.05) in *Sgcd*⁻/⁻ and wild-type Soleus muscle associated with regulation of innate immune response and response to interferon-beta pathways. Color scale represents VST-normalized expression values scaled by Z-score. Hierarchical clustering was applied to rows and columns. **D)** E) Violin plots of VST-normalized expression values for differentially expressed genes in *Sgcd*⁻/⁻ EDL muscle comparing epicatechin-treated versus vehicle-treated (DMSO) groups, corresponding to genes shown in panel C. Two-tailed unpaired Student’s t-test: *p < 0.05, **p < 0.01, ***p < 0.001. **E)** Bar plot of estimated immune cell abundance in EDL muscles from *Sgcd*⁻/⁻ vehicle-treated (DMSO), *Sgcd*⁻/⁻ epicatechin-treated, and wild-type mice, as determined by mMCP-counter deconvolution (n = 3 per group). **F)** Estimated abundance of M1 macrophages in EDL muscles from *Sgcd*⁻/⁻ vehicle-treated (DMSO), *Sgcd*⁻/⁻ epicatechin-treated, and wild-type mice, as determined by quanTIseq deconvolution. Data are presented as mean ± SD (n = 3 per group). One-way ANOVA followed by Šídák’s post hoc test: *p < 0.05, **p < 0.01, ***p < 0.001.

Since genes associated with inflammation and cytokine-cytokine communication were repressed in response to (-)-Epicatechin, we explored whether changes on inflammatory infiltrate would account for this reduction. Deconvolution analyses showed that the proportion of monocytes/macrophages diminished after treatment, whereas basophils and endothelial cells increased (Figure 3F). More importantly, the proportion of M1-like macrophages decreased, relative to vehicle-treated animals (Figure 3G; supplementary figure 6A). These results show that (-)-Epicatechin ameliorates chronic inflammation in the soleus deficient of δ-sarcoglycan.

In contrast, when we analyzed transcriptional changes in EDL after (-)-Epicatechin treatment, only subtle changes in gene expression were observed, evidenced in Figure 4A and by the low content of genes represented in GO analyses (Figure 4B; supplementary List 11). Although (-)-Epicatechin induced the reduction of genes involved in pathways such as regulation of innate immune response and cellular response to interferon beta, the number of genes and the level of reduction was not as significant as the ones observed for soleus (Figure 4C). Genes within these pathways included Gm12185, Gbp7, Gbp3, Gbp5, Trim30d, Stat1, Stat2, and Oas1a, among others (Figure 4D). In line with this, deconvolution analyses showed no differences in the transcriptional signatures of inflammatory infiltrate between (-)-Epicatechin and DMSO treatments in EDL of *Sgcd^−/-^* mice (Figure 4E). Indeed, when we analyzed the proportion of macrophages between treated and control animals, M1-like macrophages did not decrease in response to (-)-Epicatechin (Figures 4F, supplementary figure 6B), demonstrating that soleus is more responsive to the anti-inflammatory effects of (-)-Epicatechin in the context of LGMD-2F.

In contrast, (-)-Epicatechin induced a broader transcriptional response in wild-type mice than in Sgcd⁻/⁻ muscles. In wild-type Soleus, 101 genes were upregulated and 158 were downregulated (supplementary list 17), while in wild-type EDL, 272 genes were upregulated and 199 were downregulated (supplementary list 18). GO enrichment analysis revealed that upregulated genes in wild-type muscles were associated with actomyosin structure organization, muscle cell development and myofibril, whereas downregulated genes were enriched in MHC-class-II protein complex and collagen binding (Supplementary Figure 7; supplementary List 19).

In summary, our results highlight that although inflammation plays a critical role in the development of muscular dystrophy, intrinsic aspects of genotype and fiber-type may account for differential damage.

## Discussion

In the present study, we conducted a transcriptomic characterization of soleus and EDL muscles from a δ-SG-null mouse model, as well as the transcriptional changes induced by (−)-Epicatechin treatment. Our main goal was to investigate fiber-type-specific drivers of muscle damage in LGMD-2F. We found no evidence of fiber-type switching; however, structural and contraction-related genes were broadly downregulated in both muscles, indicating widespread transcriptional disruption. δ-SG-null mice exhibited a significant transcriptional shift, involving over 3,000 differentially expressed genes across muscle groups (Supplementary List 12). Notably, the enrichment of immune-related pathways—including NF-κB, Toll-like receptors, and inflammasome signaling—supports a model in which chronic inflammation is a primary driver of pathology, particularly in soleus, where inflammatory features were more pronounced. Consistently, (−)-Epicatechin reduced transcriptional markers of inflammatory infiltrate, with the strongest effect observed in soleus.

A substantial fraction of the differentially expressed genes identified in the δ-SG-null mice overlaps with those reported in other muscular dystrophies, supporting shared pathogenic mechanisms. Among these, inflammatory mediators such as Isg15, Tlr2, and Tlr4—previously linked to inflammatory myopathies and dystrophic conditions^[47–50]^—were upregulated. In parallel, and consistent with alterations described in sarcoglycanopathies^[51]^, we observed increased expression of acetylcholine receptor subunits Chrnb1, Chrng and Chrnd (Supplementary list 12), reinforcing the emerging role of the sarcoglycan complex at the neuromuscular junction. All these findings indicate that this model recapitulates key inflammatory and neuromuscular features of muscular dystrophies.

Recent evidence further supports a conserved inflammatory signature across models and patients. Amaro et al. (2025) reported increased immune infiltrate in severe LGMDR3 cases, predominantly composed of monocytes, T-cells, cytotoxic lymphocytes, and dendritic cells. A similar pattern is observed in the α-SG-null mouse model, where transcriptomic deconvolution revealed increased abundance of monocytes, T lymphocytes, and dendritic cells in quadriceps^[6]^. This convergence across human and murine systems supports a shared pro-inflammatory landscape.

Consistent with previous reports in LGMD-2F using the same δ-SG-null model, early muscle fiber damage is followed by a rapidly maladaptive inflammatory response, characterized by increased expression of pro-inflammatory cytokines (Tnfa, Il1b, Il6) and immune cell markers (Ptprc, Ly6c1, Mrc1)^[52]^. Here, analysis at a comparable disease stage revealed a similar pattern, with stronger induction in soleus. Deconvolution analyses further indicated enrichment of M1-like macrophages alongside reduced endothelial signatures, supporting the notion that early pro-inflammatory activation is an intrinsic feature of δ-sarcoglycan deficiency (Figure 2E-F). These findings align with our previous observations of preferential damage in soleus^[5]^, linking oxidative fibers to an exacerbated inflammatory phenotype.

Furthermore, metabolic properties of oxidative muscle provide a mechanistic basis for this susceptibility. Under physiological conditions, high mitochondrial density sustains efficient substrate utilization and limits inflammatory signaling^[53]^. However, sarcolemmal instability promotes calcium influx ^[54–56]^, triggering mitochondrial dysfunction marked by elevated reactive oxygen species (ROS) production and cytosolic release of mitochondrial DNA^[57–60]^. These alterations amplify inflammatory signaling via ROS and mitochondrial-derived damage-associated molecular patterns (DAMPs), leading to activation of innate immune pathways such as cGAS–STING^[31,61,62]^. Under chronic conditions, these signals sustain type-I interferon responses and converge with ROS-driven activation of the NLRP3 inflammasome and NF-κB, promoting expression of IL-1β, TNF-α, and IL-6^[25,26,63]^. Together, these mechanisms position mitochondrial dysfunction as a central amplifier of inflammation in oxidative muscles such as the soleus.

In this context, and within the same δ-SG-null model, lipid-mediated signaling may further exacerbate disease progression. Dysregulation of the autotaxin–lysophosphatidic acid (ATX–LPA) axis and activation of YAP/TAZ have been reported in triceps, gastrocnemius, and diaphragm, linking lipid signaling, mechano-transduction, and pro-fibrotic gene programs^[52]^. These findings suggest that mitochondrial dysfunction and lipid-flux may converge to potentiate inflammatory and fibrotic responses in oxidative fibers. Although soleus was not directly assessed, its pronounced susceptibility observed here highlights the need to evaluate whether these mechanisms are further amplified in this muscle.

Building on this framework, the effects of (−)-Epicatechin can be interpreted as coordinated modulation of metabolic and inflammatory networks. Previous studies have shown that (−)-Epicatechin influences mitochondrial function, redox balance, and inflammatory signaling^[44,46,64,65]^. Here, treatment reduced the M1-like macrophage signature in soleus (Figure 3G), highlighting modulation of pro-inflammatory activity as a central component of its mechanism of action. In addition, genes associated with myotonic dystrophy type-2 (DM2) and linked to cGAS–STING signaling (e.g., Irf7, Ifit3, Mx1, Ddx58/RIG-I)^[31]^ were upregulated in δ-SG-null mice and effectively downregulated following treatment, reinforcing its role in attenuating inflammation-associated transcriptional programs.

Emerging evidence suggests that flavonoids may act through receptor-mediated mechanisms, including signaling via the G protein-coupled estrogen receptor (GPER), which regulates macrophage function and inflammatory responses^[43,66]^. Although not directly assessed here, GPER provides a plausible link between extracellular signals and intracellular metabolic–inflammatory responses. Notably, GPER expression increases in inflamed skeletal muscle and correlates with disease severity^[67]^, potentially contributing to the enhanced responsiveness of soleus to (−)-Epicatechin. In this context, higher inflammatory burden may sensitize oxidative muscles to receptor-mediated signaling, supporting GPER as a potential mediator of muscle-type-specific effects. Additionally, GPER is expressed in immune cells, where it modulates inflammatory signaling through attenuation of NF-κB and reduced cytokine production^[68–70]^.

In parallel, activation of energy-sensing pathways such as AMPK by (−)-Epicatechin may further integrate metabolic and immune responses^[42,45]^. AMPK has been implicated in macrophage polarization, promoting a shift away from pro-inflammatory states toward reparative phenotypes^[41,71]^. Accordingly, the reduction in M1-like macrophage signatures observed here may reflect, at least in part, AMPK-dependent modulation of immune cell states, supporting a model in which epicatechin operates at the interface of metabolism and immune regulation.

A notable feature of our results is the muscle-type-specific response to (−)-Epicatechin. Wild-type animals showed a stronger transcriptional response in EDL, whereas δ-SG-null mice exhibited a more pronounced response in soleus. In wild-type muscle, this response was dominated by upregulation of structural genes, with minimal effects on immune pathways (Supplementary Figure 7), suggesting that pre-existing damage may be required to reveal the anti-inflammatory effects of (-)-Epicatechin. Consistently, under regenerative conditions, (-)-Epicatechin accelerated muscle recovery in wild-type mice (Supplementary Figure 5). This encourages further exploration of (-)-Epicatechin not only in muscle-wasting diseases but also in contexts with mild regenerative stimuli, such as intense exercise or minor trauma.

Finally, a limitation of this study is the lack of functional validation of immune cell profiles across muscle types and treatments. Nevertheless, our findings provide insight into fiber-type-specific susceptibility to inflammation and damage in LGMD-2F. Whether these mechanisms extend to other muscular dystrophies remains an important question for future studies and may inform the development of targeted therapeutic strategies.

## Conclusion

In summary, our findings provide experimental evidence explaining differential susceptibility to damage between muscle groups in muscular dystrophy. Transcriptomic analyses suggest that the inflammatory response in oxidative muscle is more persistent than in EDL. As a consequence, (-)-Epicatechin treatment ameliorates this response. Finally, our data suggest that this anti-inflammatory effect is more effective under muscle damage conditions.

## Acknowledgments

We thank María Luisa Benitez-Hess, PhD for technical assistance and laboratory management during the development of this project.

## Author Contributions

A.C-B. and M.H-S.: Experimental design, Methodology and Data curation and Analysis, Writing original draft. E.B-C., E.G.G-G and M.R-R.: Data analysis, Reviewing and Editing. O.H-H. and R.M.C-V.: Funding acquisition, Reviewing and Editing. J.M.H-H: Project design. Data analysis. Project administration. Funding acquisition. Conceptualization, Supervision, Reviewing and Editing.

## Data Availability Statement

For reviewing purposes, all raw data generated in this study are available under the access GSE327571, token: qzafkokqlvsrbot

## SUPPLEMENTARY INFORMATION

**Supplementary Figure 1. Profiling of global upregulated genes in skeletal muscles from *Sgcd⁻/⁻* mice compared to wild-type controls. A)** Volcano plot comparing gene expression in *Sgcd^−/-^*. **B)** Functional enrichment analysis of genes that were upregulated in *Sgcd⁻/⁻* versus wild-type mice. Differentially expressed genes were identified using DESeq2, with upregulated genes defined as those with log2FC > 0 and an adjusted p-value of ≤ 0.05. Gene Ontology (GO) enrichment analysis was performed for the Biological Process (BP), Cellular Component (CC) and Molecular Function (MF) categories. **C)** Cnetplot visualization of GO enrichment results, highlighting two representative pathways for each GO category. Lines connect individual genes to their associated enriched pathways, illustrating gene-pathway relationships. **D)** KEGG pathway enrichment analysis of upregulated genes was performed using the enrichKEGG function. The significance thresholds for all enrichment analyses were set at an adjusted p-value of <0.05 and a q-value of <0.2. The top 20 most significantly enriched terms are shown. All enrichment analyses were performed using mouse gene annotations (Mus musculus; org.Mm.eg.db for GO and organism = “mmu” for KEGG).

**Supplementary Figure 2. Profiling of global downregulated genes in skeletal muscles from *Sgcd⁻/⁻* mice compared to wild-type controls. A)** Functional enrichment analysis of genes that were downregulated in *Sgcd⁻/⁻* versus wild-type mice. Differentially expressed genes were identified using DESeq2, with upregulated genes defined as those with log2FC < 0 and an adjusted p-value of ≤ 0.05. Gene Ontology (GO) enrichment analysis was performed for the Biological Process (BP), Cellular Component (CC) and Molecular Function (MF) categories. **B)** Cnetplot visualization of GO enrichment results, highlighting two representative pathways for each GO category. Lines connect individual genes to their associated enriched pathways, illustrating gene-pathway relationships. **C)** KEGG pathway enrichment analysis of downregulated genes was performed using the enrichKEGG function. The significance thresholds for all enrichment analyses were set at an adjusted p-value of <0.05 and a q-value of <0.2. The top 20 most significantly enriched terms are shown for each analysis. All enrichment analyses were performed using mouse gene annotations (Mus musculus; org.Mm.eg.db for GO and organism = “mmu” for KEGG).

**Supplementary Figure 3. Immune response-related genes show significantly increased expression in *Sgcd^−/-^* soleus versus *Sgcd^−/-^* EDL muscles. A)** UpSet plot showing the overlap of differentially expressed genes (adjusted p-value ≤ 0.05) across four gene sets: upregulated and downregulated genes in *Sgcd*⁻/⁻ EDL and *Sgcd*⁻/⁻ Soleus muscles relative to their respective wild-type controls. Differential expression analysis was performed independently for each muscle using DESeq2. Upregulated and downregulated genes were defined as log₂FC > 0 and log₂FC < 0, respectively. **B)** Scatter plot of log₂ fold-change values for 1,700 shared DEGs (from panel A) between EDL (y-axis) and Soleus (x-axis) muscles. Each point represents a gene. The dashed line indicates the identity line (x = y). Spearman rank correlation: ρ = 0.935, p < 2.2 × 10⁻¹⁶. Red points indicate ten genes with discordant direction of regulation between tissues. **C)** RT-qPCR validation of *Tlr2*, *Tlr4*, *Myd88*, *Irf7*, and *Cxcl9*. Expression was normalized to *Tbp* and presented as a fold-change relative to *Sgcd^−/-^* EDL. Data represent mean ± SD (n = 3 biological replicates per group). Two-tailed unpaired Student’s t-test comparing *Sgcd^−/-^* Soleus versus *Sgcd^−/-^* EDL: *p < 0.05, **p < 0.01, ***p < 0.001.

**Supplementary Figure 4. *Sgcd*⁻/⁻ Soleus muscle exhibits elevated immune-related gene expression compared to wild-type. A)** Estimated abundance of ten immune cell types in Soleus and EDL muscles from wild-type and *Sgcd*⁻/⁻ mice, as determined by quanTIseq deconvolution (n = 3 per group). **B)** RT-qPCR validation of 12 immune-related genes in *Sgcd*⁻/⁻ Soleus muscle relative to wild-type controls. Expression levels were normalized to *Tbp* and presented as fold-change relative to wild-type. Data represent mean ± SD (n = 3 biological replicates per group). Two-tailed unpaired Student’s t-test: *p < 0.05, **p < 0.01, ***p < 0.001.

**Supplementary Figure 5. (-)-Epicatechin reduces tissue damage in BaCl₂-injured tibialis anterior muscle. A)** Representative transverse sections of tibialis anterior (TA) muscle stained with Gomori trichrome at 2, 4, and 15 days after BaCl₂-induced injury (1.2%). Muscle fibers appear in red; areas of tissue damage (fiber loss, inflammatory infiltrate, and interstitial space expansion) appear as pale regions. Three conditions are shown: uninjured, injured vehicle-treated and injured epicatechin-treated. Insets show lower magnification views. Scale bars: 100 μm. **B)** Quantification of damaged area relative to total muscle cross-sectional area at 2, 4, and 15 days post-injury in vehicle-treated and epicatechin-treated groups. Data represent mean ± SD (n = 3). Two-way ANOVA with Šídák’s post hoc test: **p < 0.01, ***p < 0.001; ns, not significant.

**Supplementary Figure 6. (-)-Epicatechin reduces M1 macrophage abundance in *Sgcd*⁻/⁻ Soleus but not EDL muscle. A)** Estimated abundance of ten immune cell types in Soleus muscles from wild-type, Sgcd⁻/⁻ vehicle-treated (DMSO), and Sgcd⁻/⁻ epicatechin-treated mice, as determined by quanTIseq deconvolution (n = 3 per group). **B)** Estimated abundance of ten immune cell types in EDL muscles from wild-type, Sgcd⁻/⁻ vehicle-treated (DMSO), and Sgcd⁻/⁻ epicatechin-treated mice, as determined by quanTIseq deconvolution (n = 3 per group).

**Supplementary Figure 7. (-)-Epicatechin induces distinct transcriptional responses in wild-type and *Sgcd*⁻/⁻ muscles. A)** and **B)** Volcano plots of differentially expressed genes comparing epicatechin-treated versus vehicle-treated (DMSO) wild-type mice in Soleus (A) and EDL (B) muscles. Red points indicate significant genes (adjusted p-value ≤ 0.05); gray points indicate non-significant genes. Differential expression analysis was performed using DESeq2 (n = 3 biological replicates per group). **C)** and **D)** Gene Ontology (GO) enrichment analysis of differentially expressed genes from panels A and B, respectively. C) Soleus; D) EDL. Upregulated genes (log₂FC > 0, adjusted p-value ≤ 0.05) and downregulated genes (log₂FC < 0, adjusted p-value ≤ 0.05) were analyzed separately. Biological Process (BP), Cellular Component (CC), and Molecular Function (MF) categories are shown. Enrichment analysis was performed using clusterProfiler (adjusted p-value < 0.05, q-value < 0.2). **E)** Venn diagrams showing the overlap of upregulated and downregulated genes (epicatechin-treated versus vehicle-treated) between wild-type and *Sgcd*⁻/⁻ mice in Soleus and EDL muscles. Differentially expressed genes were defined as adjusted p-value ≤ 0.05 with log₂FC > 0 (upregulated) or log₂FC < 0 (downregulated). **F)** Summary table of overlapping upregulated and downregulated genes between wild-type and *Sgcd*⁻/⁻ Soleus and EDL muscles in response to epicatechin treatment, corresponding to the Venn diagrams in panel E.

## SUPPLEMENTARY LISTS

**List_1_Fig.1C_volcano_WT.xlsx**

**List_2_Fig.1D_GOenrichment_WT.xlsx**

**List_3_Fig.2A_Volcano_Sgcdnull_SoleusvsWT_Soleus.xlsx**

**List_4_Fig.2B_Volcano_Sgcdnull_EDLvsWT_EDL.xlsx**

**List_5_Fig.2C_GOenrichment_Sgcdnull_Soleus.xlsx**

**List_6_Fig.2D_GOenrichment_Sgcdnull_EDL.xlsx**

**List_7_Fig.2G_Heatmap_comparison.xlsx**

**List_8_Fig.3B_Volcano_Sgcdnull_soleus_epicatechin.xlsx**

**List_9_Fig.3C_GOenrichment_Sgcdnull_Soleus_Epicatechin.xlsx**

**List_10_Fig.4A_Volcano_Sgcdnull_EDL_epicatechin.xlsx**

**List_11_Fig.4B_GOenrichment_Sgcdnull_EDL_Epicatechin.xlsx**

**List_12_Fig.Sup1A_Volcano_sgcdnull_vs_WT.xlsx**

**List_13_Fig.Sup1B_GOenrichement_Sgcdnull_UP.xlsx**

**List_14_SupFig1D_KEGG_sgcdnull_vs_WT_UP.xlsx**

**List_15_SupFig2A_GOenrichment_Sgcdnull_Down.xlsx**

**List_16_SupFig2C_KEGG_sgcdnull_vs_WT_Down.xlsx**

**List_17_SupFig7A_WT_soleus_epicatechin.xlsx**

**List_18_SupFig7B_WT_EDL_epicatechin.xlsx**

**List_19_SupFig7C-D_GOenrichment_epicatechin.xlsx**

---

**Supplementary table 1.**
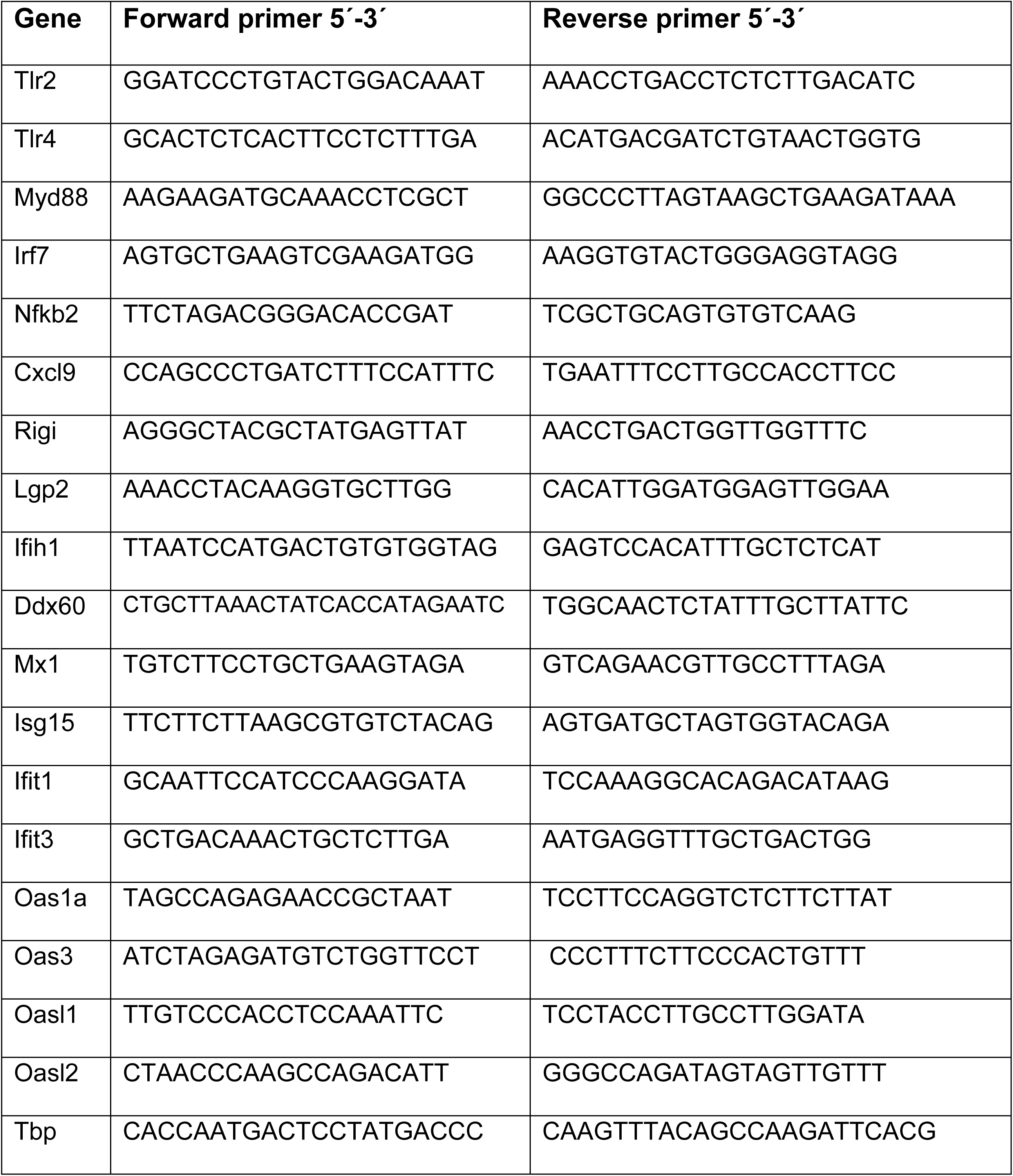
Primers used for the target genes in this study.

